# Automated quantification of lipophagy in Saccharomyces cerevisiae from fluorescence and cryo-soft X-ray microscopy data using deep learning

**DOI:** 10.1101/2023.02.27.530171

**Authors:** Jacob Marcus Egebjerg, Maria Szomek, Katja Thaysen, Alice Dupont Juhl, Stephan Werner, Christoph Pratsch, Gerd Schneider, Richard Röttger, Daniel Wüstner

**Affiliations:** Department of Biochemistry and Molecular Biology, University of Southern Denmark, Campusvej 55, DK-5230 Odense M, Denmark; Department of Mathematics and Computer Science, University of Southern Denmark, Campusvej 55, DK-5230 Odense M, Denmark; Department of X-Ray Microscopy, Helmholtz-Zentrum Berlin, Albert-Einstein-Str. 15, 12489 Berlin, Germany and Humboldt-Universität zu Berlin, Institut für Physik, 12489 Berlin, Germany

**Keywords:** deep learning, extracellular vesicles, fluorescence, intracellular, lipophagy, lipid droplets, phenotype, segmentation, X-ray microscopy, yeast

## Abstract

Lipophagy is a form of autophagy by which lipid droplets (LDs) become digested to provide nutrients as a cellular response to starvation. Lipophagy is often studied in yeast, *Saccharomyces cerevisiae*, in which LDs become internalized into the vacuole. There is a lack of tools to quantitatively assess lipophagy in intact cells with high resolution and throughput. Here, we combine soft X-ray tomography (SXT) with fluorescence microscopy and use a deep learning computational approach to visualize and quantify lipophagy in yeast. We focus on yeast homologs of mammalian Niemann Pick type C proteins, whose dysfunction leads to Niemann Pick type C disease in humans, i.e., NPC1 (named NCR1 in yeast) and NPC2. We developed a convolutional neural network (CNN) model which classifies ring-shaped versus lipid-filled or fragmented vacuoles containing ingested LDs in fluorescence images from wild-type yeast and from cells lacking NCR1 (*Δncr1* cells) or NPC2 (*Δnpc2* cells). Using a second CNN model, which performs automated segmentation of LDs and vacuoles from high-resolution reconstructions of X-ray tomograms, we can obtain 3D renderings of LDs inside and outside of the vacuole in a fully automated manner and additionally measure droplet volume, number, and distribution. We find that cells lacking functional NPC proteins can ingest LDs into vacuoles normally but show compromised degradation of LDs and accumulation of lipid vesicles inside vacuoles. This phenotype is most severe in *Δnpc2* cells. Our new method is versatile and allows for automated high-throughput 3D visualization and quantification of lipophagy in intact cells.

## Introduction

Lipids are important for the structural integrity of cellular membranes and also function as regulators of membrane fusion and fission events. Beside membrane lipids, cells store triacyl esters of glycerol and fatty acyl esters of sterols as neutral lipids in lipid droplets (LDs). Lipophagy is a cellular process that involves the degradation of LDs within a cell through autophagy. It is a mechanism by which cells can clear excess lipids and prevent lipid accumulation, playing an important role in maintaining cellular energy homeostasis and regulating lipid metabolism. Lipophagy is gaining interest as an important cellular pathway, not only for mobilizing stored lipids and providing precursors for membrane synthesis, but also as potential mechanism for clearance of miss-folded or aggregated proteins, which have been shown to accumulate on the droplet surface [1, 2]. Defective lipid trafficking is a hallmark of many neurodegenerative diseases, including Niemann Pick type C (NPC) disease. In this lipid storage disorder mutations in either Niemann Pick type C1 or C2 proteins (NPC1 and NPC2) lead to accumulation of cholesterol and sphingolipids, such as sphingosine, glycosylceramide and sphingomyelin, in lysosomes [3, 4]. Mammalian NPC2 has been proposed to pick up cholesterol derived from low density lipoprotein and other sources inside lysosomes and deliver it to NPC1 for integration into the lysosomal membrane [5, 6]. Recently, both proteins have been implicated in autophagy in yeast, as normal vacuole function and lipid mobilization from LDs were compromised in cells lacking functional NPC proteins during starvation [7, 8]. Yeast vacuoles retain many functions of mammalian lysosomes, and they are due to their larger size amenable to microscopic observation. This, together with the much simpler genetic manipulation compared to mammalian cells make yeast cells a widely established model for studying lipophagy and other autophagic processes [2, 9]. Also, NCR1, the yeast homolog of NPC1, and yeast NPC2 show high structural similarity to mammalian NPC proteins, and both, NPC1- and NPC2-deficient mamalian cells can be rescued by the *S. cerevisiae* counterpart [10, 11].

In many cell biological studies lipophagy is assessed by staining vacuoles and LDs using suitable and complementary dyes, such as FM4-64 and BODIPY 493/503, respectively. The number of ingested LDs is typically assessed by manual counting in epifluorescence or confocal images, a tedious procedure, which suffers from the limited resolution of conventional fluorescence microscopy which at best allows one to discern cellular sturctures being about 300 nm apart. Furthermore, more subtle phenotypes might be missed by this approach, for example strucural changes in vacuoles or accumulation of other lipid assemblies inside vacuoles apart from LDs. To improve the resolution, various approaches have been used; e.g., single molecule localization microscopy of BODIPY dyes or photoactivatable fluorescent proteins can provide information about droplet localization inside and outside of vacuoles with a resolution of 30-50 nm [12, 13]. This approach, however, does not alleviate the need for staining, thereby largely missing the cellular context in which lipophagy and other autophagic processes take place. Alternatively, electron microscopy (EM) can provide ultrastructural information of vacuoles and LDs with < 10-nm resolution and in a label-free manner but only for thin slices due to the low penetration depth of electrons in biological samples [14]. Freeze-etch EM is based on fracturing frozen specimen, thereby obtaining a replica of the original biological structure [15]. Depending on the used freezing technique and fracture protocol, membrane deformation and structural rearrangements cannot be excluded in this technique [16]. The best possible resolution is currently obtained with focused ion beam milling scanning electron microscopy (FIB-SEM), allowing for attaining close-to molecular resolution in three dimensions. This comes to the price of slow acquisition speed and need for proper fixation and embedding of the specimen with an average acquisition time of 2-5 days per cell [17]. Soft X-ray tomography (SXT) is a relatively new imaging method which achieves 3D information with isotropic 30-nm resolution of intact cryo-frozen cells throughout their entire thickness [18, 19]. Its contrast is based on the extent by which organelles absorb X-rays which is high for lipid membranes and LDs and low for the cytosol or water-filled vacuole, allowing for ultrastructural analysis of cells including starvation responses in yeast [20, 21, 22, 23]. SXT has an about 10-fold higher resolution than diffraction-limited fluorescence microscopy but about 5-fold lower resolution compared to FIB-SEM. However, SXT is much faster than FIB-SEM as it allows for 3D imaging of entire cells in their near-native state in about half an hour. Also, SXT is not invasive and does not require cell fixation, in contrast to FIB-SEM. SXT relies on the energy range known as the *water window*, which is defined as the X-ray absorption of the *K* edges of oxygen (543 eV, 2.28 nm) and carbon (284 eV, 4.37 nm) [24]. By working within this energy window there is no need for additionally staining of biological samples, hence a natural absorption contrast will emerge from the biological carbon-rich structures. SXT is therefore ideally suited to study cellular ultrastructure at the mesoscale, bridging the gap between electron and light microscopy. SXT can be combined with cryo fluorescence microscopy, allowing for subcellular imaging and phenotype mapping with much higher throughput than possible in EM. A bottleneck in the analysis of 3D image stacks reconstructed from X-ray tomograms is the lack of suitable software tools for organelle segmentation, as only very few studies have attempted to use modern machine learning approaches to segment X-ray images in an automated manner for selected problems [25, 26]. This leaves the segmentation task often to tedious manual annotation of images in specialized settings and research environments.

In this study, we employ a combination of SXT and fluorescence microscopy and use deep learning to quantify the process of lipophagy in yeast cells. To accomplish this, we have created two convolutional neural network (CNN) models; the first model is used for automated classification of yeast vacuoles at different stages of lipophagy from fluorescence images. The second CNN model allows for automated segmentation of X-ray images and thereby to measure the ingestion of LDs into vacuoles of both wild-type yeast and yeast cells lacking either NCR1 or NPC2 from X-ray tomograms. By this computational approach we are able to quantify impaired lipophagy and compromised vacuole morphology and to create fully automated 3D renderings of LDs inside and outside of the vacuole to measure droplet volume, number, and distribution. We discovered that yeast cells lacking functional NPC proteins have partially fragmented vacuoles and a compromised ability to degrade ingested LDs inside the vacuoles. Our novel combination of automated phenotype classification and instance segmentation based on combined fluorescence and X-ray microscopy data is versatile and provides a high-throughput method for 3D visualization and quantification of lipophagy in intact cells. Combined with genetic knock-down and functional experiments, our method will provide mechanistic inside into lipophagy in various physiological settings.

## Results

### A convolutional neural network to classify vacuolar phenotypes from fluorescence images

When yeast cells are kept in the same culture medium during stationary growth, they become depleted of nutrients, resulting in several starvation responses, including lipophagy, the ingestion of LDs rich in sterol and triacylglycerol esters into the vacuole [2]. We have shown previously using fluorescence microscopy, that NCR1 and NPC2 are instrumental for transport of sterols into the vacuole during stationary growth [8]. Absence of functional NPC proteins diminished incorporation of the fluorescent ergosterol analogue dehydroergosterol (DHE) into the vacuolar membrane, which was labeled with the lipophilic dye FM4-64 (Figure 1A and B). While DHE is inserted into the vacuole membrane in wild-type cells, it accumulates instead in LDs stained with BODIPY 493/503 in *Δncr1* cells (Figure 1B) and *npc2* cells (not shown). The vacuole often appeared lipid-filled with diffuse intravacuolar staining with FM4-64 and some vacuoles were fragmented/multivesicular in cells lacking functional NCR1 or NPC2 [8]. This is in contrast to wild-type cells, where one central, ring-shaped vacuole was observed (Figure 1A), as being characteristic for yeast cells in the stationary phase [27]. In addition, lipid-filled vacuoles accumulated strong fluorescence of BODIPY493/503 in droplets in contrast to control cells (Figure 1A and B). Phenotypes of defective vacuolar ingestion of sterol-rich LDs in NPC-deficient yeast were previously assessed based on visual inspection and manual counting similar as in other studies on lipophagy in yeast [7, 8, 14, 28]. To automatize this tedious analysis, we developed here a classification CNN which should distinguish these two phenotypes: fused vacuoles, forming a central ring in the cell with internalized and small LDs were considered as the ‘healthy’ phenotype, while multivesicular vacuoles or vacuoles filled with FM4-64 dye due to disposition of intravacuolar vesicles and LDs were categorized as ‘sick’ phenotype. The latter is likely a consequence of build-up of autophagic vesicles inside of vacuoles, since it has been shown that FM4-64 can slowly diffuse into the vacuole and label intraluminal membrane structures, including autophagic bodies [29]. A portion of cells could not be annotated into those two categories, for which a third category, named ‘unknown’, was generated. Training and validation data was manually annotated into three categories: 1) healthy cells (n = 323), 2) sick cells (n = 354) and 3) unknown (n = 109 cells), as shown in Tab. 1.

**Figure 1.**
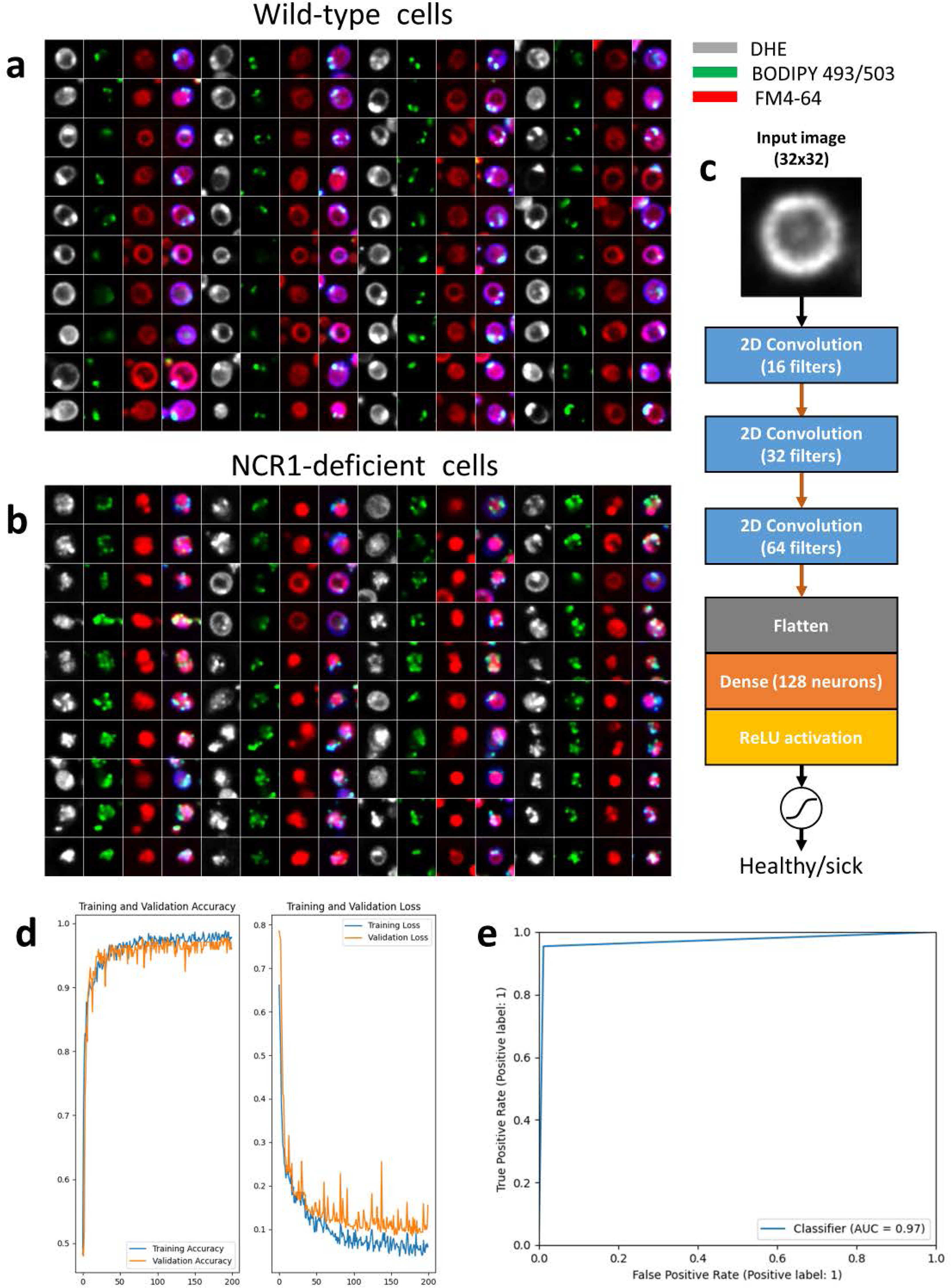
Deep learning-based classification of lipophagy phenotypes from multicolor fluorescence images. Wild-type, *Δncr1* and *Δnpc2* cells were labeled with the ergosterol analog DHE under anerobic conditions in the presence of the vacuole marker FM4-64 over night, washed and labeled with BODIPY493/503 to stain LDs for 10 min, washed and imaged. (**A and B**) Example montages of wild-type (**A**) and *Δnpc2* cells (**B**) with DHE in grey, BODIPY493/503 in green, FM4-64 in red followed by a color overlay. Four columns and ten rows giving forty cells in total are shown for each condition in a representation as see by the model. Comparable cell phenotypes as shown for *Δnpc2* cells were also found for *Δncr1* cells (not shown) but see [8]. For training and validation, only the red channel was used. (**C**) Architecture of the classification CNN using 32×32 images as input. Three blocks of repeated convolution layers for feature extraction are intervened by 2×2 max-pooling layers (red arrows) followed by a flattening layer, a dense layer and a ReLU activation layer. A sigmoid loss activation function is used for binary decision of ‘healthy’ versus ‘sick’ phenotypes. (**D**) Accuracy (left) and loss (right) for training (blue) and validation data (orange) as function of epochs. (**E**) receiver operating characteristics curve for model performance shows the ability of the classification CNN to separate both classes (AUC=area under curve). See main text for further details.

**Table 1.** Number of training and validation samples for the classification CNN.

|  | Healthy vacuole | Sick vacuole | Unknown morphology |
| --- | --- | --- | --- |
| Training | 238 | 266 | 87 |
| Validation | 85 | 88 | 22 |

The sick vacuole phenotype indicates either a lipid-filled vacuole resulting in diffuse staining with FM4-64 or a multivesicular vacuole. While the latter is difficult to discern in wide field images, we found both phenotypes by confocal microscopy (not shown but see Fig. S6 in [8]). To provide input data for the model, regions of interest of size 32×32 capturing single cells were automatically generated and extracted from all channels based on the brightfield image by performing circular Hough transformation to locate the cells in each image field (Fig. S1). The unknown category was added because the circular Hough transformation also yielded images that are not centered, contained damaged cells or cells, not being in focus or only partially captured by the segmentation. A CNN architecture was implemented consisting of 3 convolution and max pooling layers followed by a layer of densely connected neurons (Figure 1C). Convolutional and dense layers are followed by a rectifier linear unit (ReLU) for activation. The output layer is a single neuron with sigmoid activation for binary classification. Two binary classifiers were trained on the architecture – one for healthy vs sick and one for cell vs unknown. The healthy vs sick classifier achieved >97% accuracy on the validation set, indicating that the model is robust (Figure 1D). This is also reflected by the area under the curve in the receiver operating characteristic curve shown in Figure 1E. We found 75.46% of total (n=1096) and 76.47% of total vacuoles (n=1118) classified as being sick, i.e., lipid-filled or fragmented, in *Δncr1* cells and *npc2* cells, respectively. This is much more than found for control cells, where only 47.20% of all vacuoles (n=714) appeared lipid-filled or multi-vesicular. This result shows that yeast lacking functional NCR1 or NPC2 have a compromised ability for clearance of intraluminal autophagic vesicles and for vacuolar fusion, in line with our previous results [7].

### Deep learning-based segmentation of LDs and vacuoles using fluorescence and X-ray image data

To distinguish lipid-filled vacuoles from those being multivesicular, FM4-64 staining detected by wide field fluorescence microscopy is not sufficient, and much higher resolution is needed. We employed previously confocal microscopy for this task, as it has better resolution particularly along the optical axis [8]. Still, confocal imaging is diffraction-limited, and staining with FM4-64 can only report about the distribution of labeled membranes without providing a structural context. Here, we find that X-ray microscopy is ideal for this task, as it provides information about the cellular ultrastructure with an isotropic resolution of about 30 nm throughout the entire cellular volume for many cells in a reasonable amount of time. X-ray contrast is high for lipid membranes allowing for a detailed analysis of lipophagy. To acquire the 3D tomograms, the sample is placed on a special holder that is tilted around a rotation axis. The raw tilt series is subsequently aligned from fiducial markers, and 3D images of cells are reconstructed from the aligned tomograms. Yeast cells can be additionally labeled using suitable fluorescent organelle markers, and fluorescence images of such markers can be acquired in parallel to the X-ray tomograms on the same set up [24, 30, 31]. We employ this strategy to identify LDs using the droplet marker BODIPY 493/503 (emitting in green) and the vacuole marker FM4-64 (emitting in red). A typical workflow for preparing and imaging yeast cells by combined fluorescence and SXT is shown in Fig. S2.

Wild-type yeast have typically one large vacuole under starvation conditions to which LDs can dock before being ingested (Figure 2A, upper panel). In addition, we find intraluminal vesicles and small droplet-like structures inside the vacuole of wild-type cells, but mostly in relatively low abundance. In cells lacking functional NCR1 more lipid material is deposited inside vacuoles, while in *Δnpc2* yeast cells, lacking functional NPC2 protein, the vacuole appears additionally deformed in many cells, as seen by SXT (Figure 2A, middle and lower panel). LDs appear as round and particularly dark spheres in SXT, as they efficiently absorb X-ray radiation, due to their high carbon and low water content, giving them a high linear absorption coefficient [20, 22]. Differences in X-ray absorption and morphological characteristics are necessary conditions for organelle annotation, but they are often not sufficient criteria to unequivocally annotate subcellular structures in X-ray microscopy. For that reason, we co-labeled cells with FM4-64 and BODIPY493/503, allowing for unequivocal identification of vacuoles and LDs in both imaging modalities (Figure 2B). To enrich the training data for the segmentation CNN model (see below), the input frames are augmented using random rotations and elastic deformations as described by Simard et al. (2003) [32]. This method generates variations of the original X-ray images, thereby greatly reducing the burden of manual image annotation for generation of training data (Figure 2C and below). In some of the NPC-deficient cells, aberrant vacuole morphologies were also seen by SXT. This is illustrated for *Δnpc2* yeast cells, in which vacuole staining by FM4-64 is found in convoluted structures, which are clearly visible at the ultrastructural level by SXT (Figure 2D and E). LDs are found inside and attached to those fragmented vacuoles in *Δnpc2* cells while intense staining of LDs with BODIPY493/503 is found in the same region. Similar results were found for some *Δncr1* cells (not shown). The majority of vacuoles in *Δncr1* and *Δnpc2* yeast cells consisted of one large vesicle similar to control cells but with extensive intraluminal membrane accumulation. The intravacuolar lipid enrichment causes the extensive staining of FM4-64 over the entire vacuole lumen in wide field fluorescence images, which we annotated as ‘sick’ phenotype in our classification CNN model. Together, these results show the potential of combined SXT and fluorescence microscopy to study the phenotype of yeast vacuoles in control and NPC-deficient cells. It also allows us to unequivocally annotate LDs and vacuoles from X-ray images to generate trainings data for deep learning-based image segmentation.

**Figure 2.**
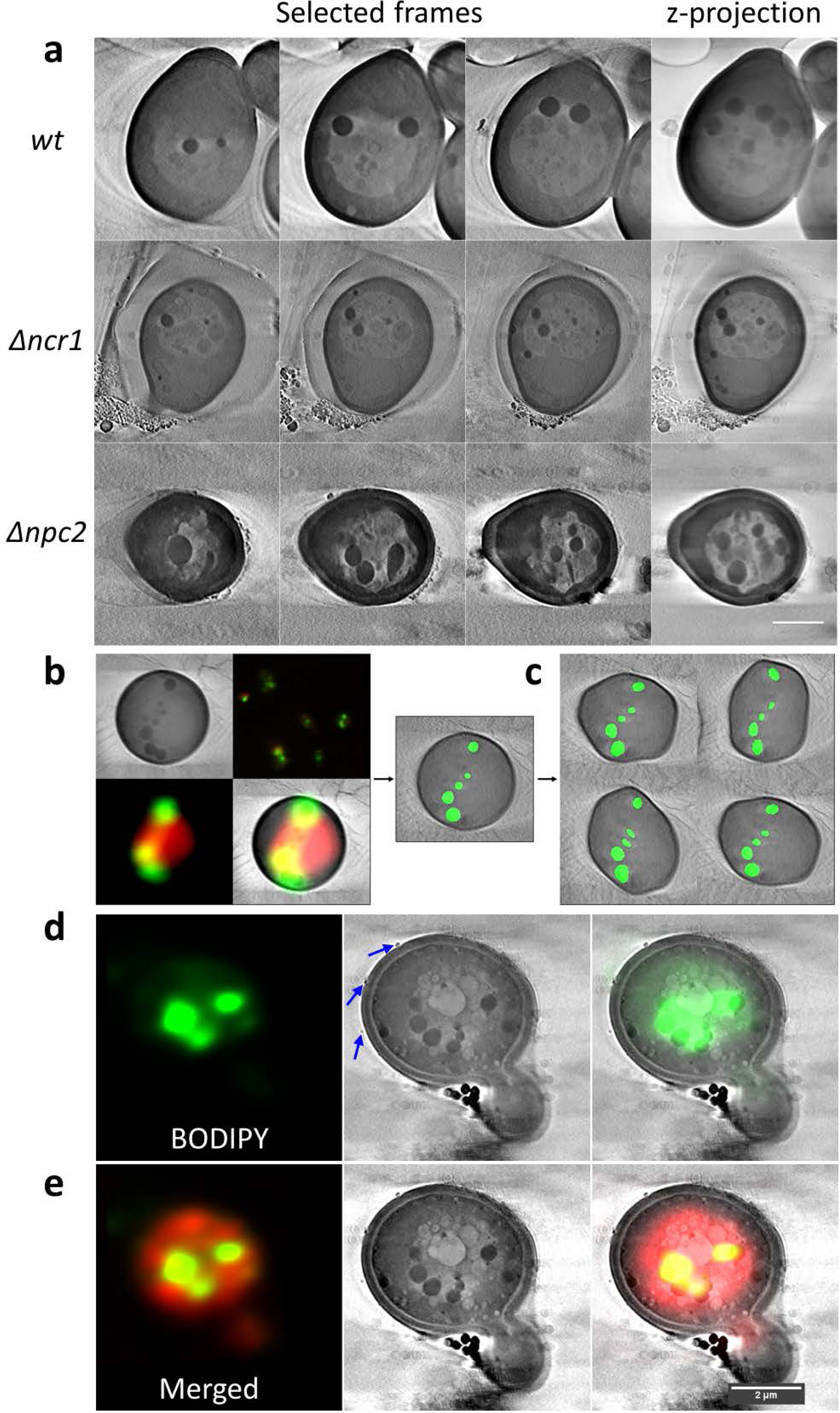
Correlative X-ray and fluorescence microscopy of wild-type and NPC-deficient yeast cells. **(A**) Selected frames (left) and z-projections (right) of SXT reconstructions for wild-type (upper row), *Δncr1* (middle row) and *Δnpc2* cells (lower row) are shown. (**B**) Correlative fluorescence and X-ray microscopy exemplified for a wild-type cell; upper left shows X-ray image, upper right, fluorescence image in original size, lower left is zoomed fluorescence image with BODIPY493/503 in green and FM4-64 in red and lower right is a color overlay. LDs are identified by the large X-ray absorption and in parallel by their green fluorescence from the BODIPY493/503 staining. (**C**) For model training, the overlayed data is augmented by non-elastic deformations to enrich the trainings data for the segmentation CNN. (**D and E**) Correlative fluorescence and X-ray microscopy of *Δnpc2* cells with a fragmented, i.e., multivesicular vacuole. Co-staining of X-ray image (middle panels) with BODIPY493/503 (**D**) and additionally with FM4-64 (**E**) is shown. Right panels are overlays of fluorescence and X-ray images.

To characterize the NPC-phenotypes at the ultrastructural level further and potentially discover more subtle differences to control cells, a pipeline was created for instance segmentation, rendering and analysis of the X-ray images (Figure 3). The first step of the pipeline is comprised of a quantile normalization to achieve comparable image histograms of the reconstructed X-ray image stacks (Figure 3A). The input for the model consists of five sequential frames from a z-stack to aid in consistency across each tomogram reconstruction, thereby alleviating issues caused by the missing wedge problem (Figure 3B). The architecture of the segmentation CNN is largely based on the original U-Net with a few changes (Figure 3C) [33, 34]. The output is a single convolutional filter with a sigmoid activation; hence, each channel is trained separately with its own set of weights. This results in a modular pipeline and simplifies the training process as each channel can have its own subset of training data and model architecture. Additionally, trained channels can be used as pre-training for other channels, which can reduce the required training time. Models were trained for cell membrane, vacuole and droplet segmentation. During training, a compound loss function consisting of Dice and TopK binary cross-entropy is used, with the K parameter available as a user parameter in the pipeline. Each training/label pair is sampled equally with respect to the biological sample, to prevent overfitting. Output of this model are the predictions for each channel (Figure 3D).

**Figure 3.**
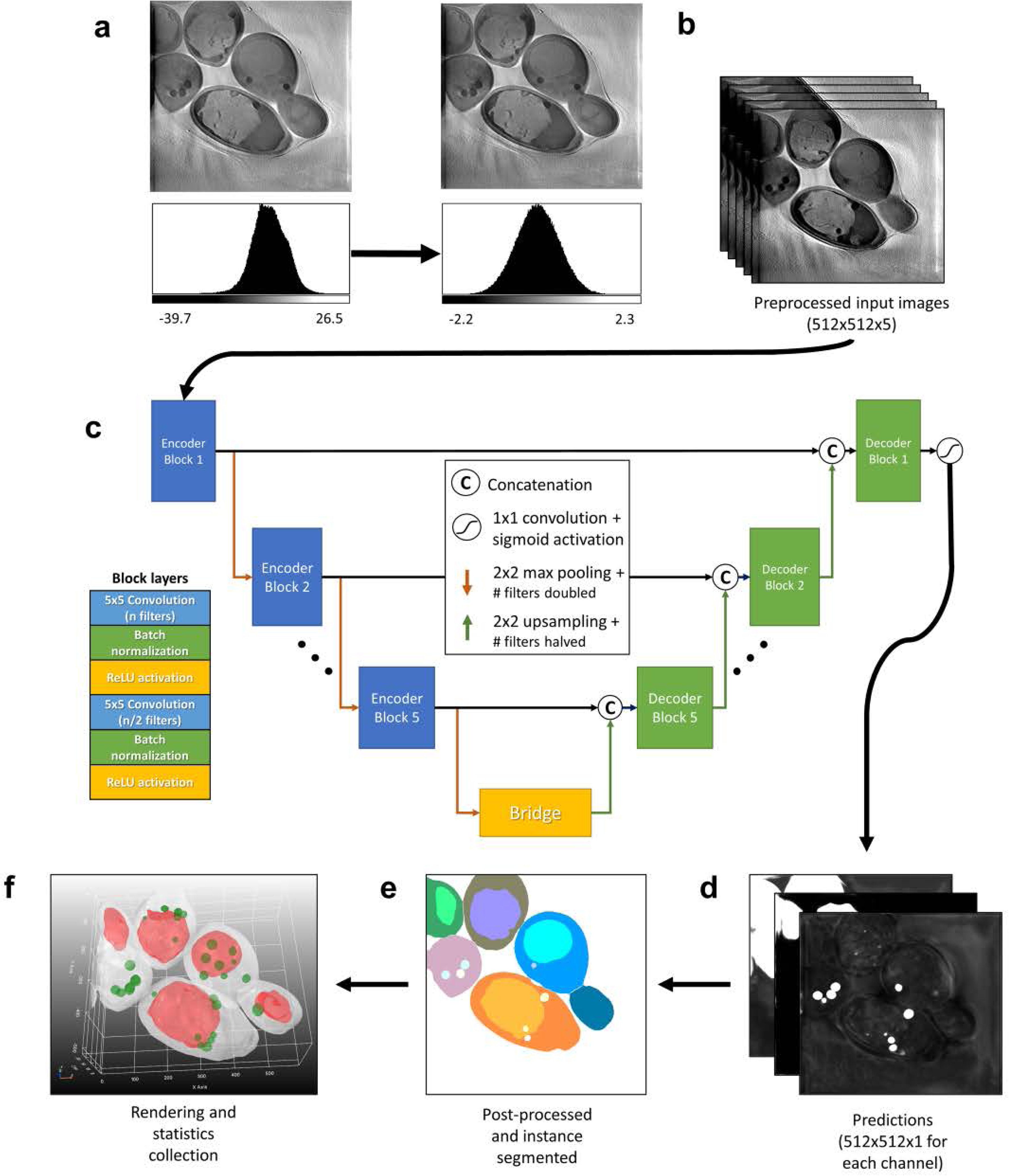
Workflow for automated segmentation and architecture of the segmentation CNN. (**A**) To obtain comparable intensity statistics for all image data sets, tomogram reconstructions were first normalized using quantile normalization based on a Gaussian distribution. (**B**) For model training, five sequential input slices were used for each stack of the training data and augmented by 90-degree rotations and elastic deformations. (**C**) The segmentation CNN model is based on the U-net architecture but with five sequential frames from a z-stack, which we found to minimize problems caused by the missing wedge problem. Each block consists of two iterations of convolution layers, batch normalization and ReLu activation unit, and blocks are connected by 2×2 max pooling layers during the encoding phase (red arrows), and 2×2 up-sampling layers during the decoding phase (green arrows). A sigmoid activation function is used as output, and each channel, i.e., entire cells, vacuoles and LDs, is trained as a separate model. During training, a compound loss function consisting of Dice and TopK binary cross-entropy is employed. (**D**) Raw predictions for each channel are generated with probability of belonging to a given class, e.g., LDs, shown in grey scale intensity values. This is followed by instance segmentation using a watershed transformation for all three channels separately (**E**). The segmented stacks are then used for 3D rendering using a marching cube algorithm for mesh generation (**F**). This is shown here as top view of model output after 3D rendering showing cells in white, vacuoles in red and LDs in green exemplified for *Δnpc2* cells. Finally, the segmented 3D objects (LDs and vacuoles) are analyzed including calculation of volumes, distances etc. See main text for further details.

To ensure robustness on a model trained on a smaller training dataset, optional post-processing steps were added. The options are outlier removal, masking, averaging/blurring, hole filling, convex constraints, minimum volume filtering and surface smoothing. Combinations of these options were used to generate new training data from unseen data, and many options are recommended, even on a well-trained model [34, 35, 36]. Given that the accuracy of the segmentation has been convincing, it discharged us from creating further manual segmentations. During development and training data creation, we tested an earlier version of the models using unseen, and partially seen test data, which has since been added to the training data. We calculated the Dice similarity coefficient (DSC) for each frame over two fully hand-segmented z-stacks (see Materials and methods) [37]. We observe a DSC of >0.9 in most frames for the cell membrane segmentations, with some exceptions in the first and last frames, where the missing wedge problem leaves the segmentation ambiguous (Fig. S3A, B, black curves). The same pattern is observed in the droplet channel, in which the DSC oscillates for the same reason (Fig. S3A, B, green curves). In most cases, the prediction of the model fits better visually, than the hand-annotated training data, explaining many of the misclassifications (Fig. S3C and D). The good agreement between X-ray images and model prediction can also be seen in the Supplemental video.

Instance segmentation is conducted using a distance transform watershed based on the implementation in MorphoLibJ, which implements a workflow using Meyer and Beucher’s watershed algorithm [38, 39]. Watershed was favored over a machine learning approach using bounding boxes or separate neural networks, such as Mask R-CNN used by Li et al. (2022) [26], to minimize constraints and workload on training data. We found that our approach was sufficient for yeast cell images, where the visible cellular morphology generally resembles ellipsoids.

A mesh is generated from the instance segmentation using marching cubes and optionally smoothened (Figure 3E and F). The meshes are rendered in an interactive 3D environment using PyVista [40]. Each mesh is saved for future rendering or analysis. Analysis could include measurement of volume, distance to nearest droplet, vacuole or cell membrane and containment of other meshes (i.e., whether LDs have been consumed by a vacuole). A representative 3D rendered image of NPC2-deficient yeast cells is shown in Figure 3F. Our model allows for accurate segmentation and 3D rendering of vacuoles and LDs in the majority of cells in a fully automated manner. Note that fragmented vacuoles or vacuoles in cells located at the edge of the tomograms could not be identified by our method. As a result, in some cells, LDs but no vacuole could be detected, while occasionally some cells had no visible droplet structure (see examples in left two cells of Figure 3F). Such cells at the edge of the images were discarded from further analysis. In most cases, however, both, vacuoles and LDs were identified and 3D-rendered and appear well-separated from each other in the 3D representation (Figure 3F, central two and right two cells).

The size of vacuoles is tightly regulated and appears largest during starvation due to pronounced homotypic vacuole fusion under this condition [41, 42]. Lipophagy starts by docking of LDs at distinct sites of the vacuole surface, and those domains are supposed to be highly ordered, rich in ergosterol and certain proteins [14, 43]. Lack of NCR1 or NPC2 was suggested to lower formation of such vacuolar domains, thereby diminishing lipophagy [7], but this could not be confirmed in another study using acute glucose restriction [44]. Both, blocking ergosterol synthesis and ergosterol overloading by preventing its esterification, seem to prevent formation of vacuolar domains, suggesting that a fine sterol balance is needed to maintain the correct vacuolaor ultrastructure during starvation [44, 45]. Based on these interesting but somehow contradicting observations, a coherent picture of vacuolar phenotypes of NPC-deficient yeast cells is lacking. Our model allows for 3D visualization and assessment of vacuolar structure and the spatial arrangement of LDs in wild-type and mutant cells, as shown in Figure 4A-F. For this, no manual segmentation or correction is needed, but 3D renderings can be obtained fully automatically with only X-ray tomogram reconstructions as input. Cells and their vacuoles appear sometimes cut off at the top or bottom as seen in side-views of 3D renderings. This is an artifact of the limited range of recorded tomographic projections, which is between +/- 80 degree but due to the ice thickness of the cryo-frozen sample maximally from −65 to +65 degrees at the BESSYII facility (see Materials and methods). Ideally, one would like to aqcuire a full-range tomogram, which requires acquistion from −90 to + 90 degrees, however, this is impossible with the current sample holder and given hardware set up. The reduced number of projections compared to a full tomogram leads to the so-called missing wedge or missing cone, a well-known problem in computerized tomography, for which currently no satisfysing algorithmic solution exists. The problem is illustrated in 2D for a simulated yeast phantom image for the filtered back projection algorithm used in this study (Fig. S4). One finds that the reduced number of projection angles, or missing wedge, causes shading artefacts at the edge of the simulated LDs, vacuoles and cell borders. The severity of such artifacts depends on the orientation of the object in the simulated image relative to the projection axis and on the distance from the image center. As a consequence, these artefacts affect the segmentation quality to varying degrees, depending on the position and orientation of the yeast cells in the ice of the cryo-sample relative to the radiation beam. The diminishing contrast at the edge of the vacuole, for example, can result in segmentation errors and cutting of the object, as seen in some 3D reconstructions. Whenever such artifacts occur in our SXT data (see an example in Figure 4D, white arrows), the vacuoles appear cut at the top and bottom. The same problem can also occasionally result in some mismatch between subjectively assessed manual segmentation of LDs and model output (Fig. S3).

**Figure 4.**
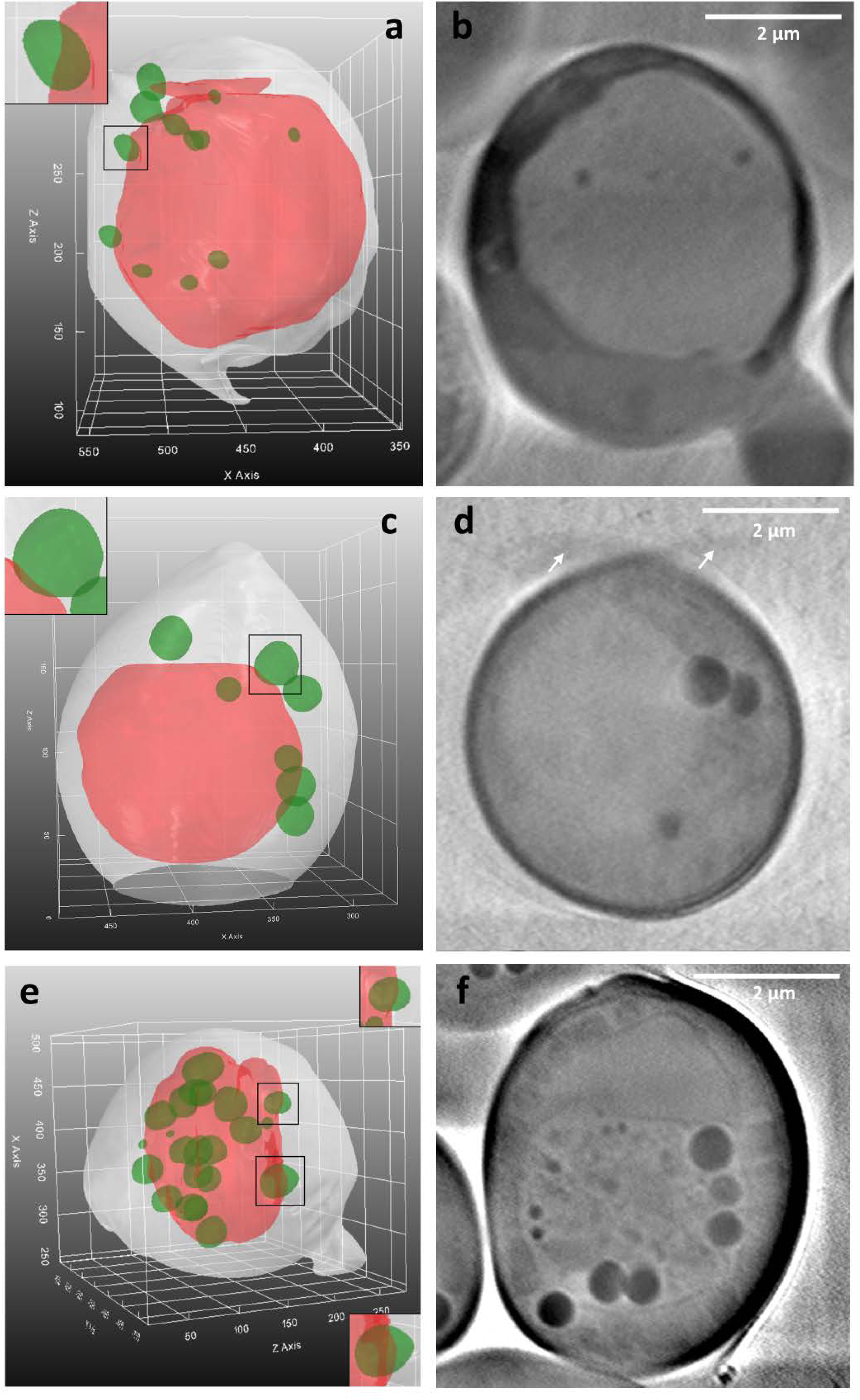
Vacuolar shapes from 3D renderings for wild-type and NPC-deficient cells. 3D renderings as obtained from the segmentation CNN model for wild-type cells (**A**), *Δncr1* cells (**C**) and *Δnpc2* cells (**E**). Insets show zoomed versions of rectangular black boxes highlighting green LDs docking on the surface of the vacuole, shown in red. Corresponding *n* sum projections for X-ray reconstructions from corresponding tomograms for wild-type cells (**B**, *n* = 20), *Δncr1* cells (**D**, *n* = 11) and *Δnpc2* cells (**F**, *n* = 5). See main text for further details.

For wild-type cells, as well as for *Δncr1* and *Δnpc2* cells, we observed docking of LDs to vacuoles, resulting in slight indentation of the vacuole membrane at the docking site (Figure 4A, C, E, insets). Such indentations have been observed previously by freeze-etch EM, and based on fluorescent markers in confocal microscopy, and those regions have been suggested to be enriched in certain proteins and lipids, necessary for droplet docking and ingestion [7, 14, 45]. The fact that we observed docking sites for LDs on vacuoles in wild-type and NPC-deficient cells with similar frequency argues against defects in the initial stage of lipophagy in cells lacking functional NCR1 or NPC2. From the 3D renderings, one can also observe the pseudo-hexagonal structure of the vacuoles, which is characteristic for the micron-scale domains found in vacuoles of starved cells [7, 14, 45, 46]. This can be clearly discerned in side-views of 3D rendering, especially for wild-type and NCR1-deficient cells. In *Δnpc2* cells we also find hexagonal-like vacuolar shapes but often additionally crinkles and an overall buckled structure (Figure 4E and F and Fig. S5). In addition, vesicles and tubular membrane structures were found inside vacuoles, and those were particularly pronounced in *Δnpc2* cells (Figure 2A and 4F and Fig. S5A-E) but also prominent in some *Δncr1* cells (Figure 2A and Fig. S5F). In some *Δnpc2* cells, we observed LDs inside of intraluminal vesicles, as previously concluded from thin EM sections [14]. While the function of such vesicles is unknown, they might protect LDs from hydrolysis by intravacuolar hydrolases or they could be the result of a different autophagic mechanism, as previously suggested [14, 47]. Most of these intravacuolar membrane assemblies are of low X-ray contrast, and this together with their heterogenous shape make them difficult to identify in an automated manner. They have therefore not been considered in our segmentation algorithm.

We also found, especially in *Δnpc2* cells, tiny vesicles at the cell wall, which are likely secreted from the cells (Figure 2D, E and Fig. S5, blue arrows). Such extracellular vesicles (EVs) have been observed previously by EM, purified from cells, and analyzed by light scattering and cryo EM tomography [48, 49]. The EVs, we find occasionally in *Δnpc2* cells by SXT are in the size range of 60-100 nm, which is in line with previous reports on EVs in yeast cells [50]. Such vesicles could be involved in cellular efflux of excess lipids, release of exosomes and autophagic membranes in yeast, as recently proposed [51]. Interestingly, using combined fluorescence and X-ray microscopy, we recently observed and characterized shedding of EVs from the plasma membrane of fibroblasts from NPC2 patients, and we showed that such vesicles contain abundant cholesterol and lysosomal cargo [30]. Thus, for both, yeast and mammalian cells, the high resolution provided by SXT allows one to study secretion of EVs. However, the rare occurrence of EVs in X-ray images of yeast makes a quantitative analysis difficult. Together, we conclude from this data, that yeast cells lacking functional NPC2 have a more severe phenotype than NCR1-deficient cells.

Using our deep learning pipeline, we are able to quantify the extent of lipophagy from X-ray reconstructions. For that, the meshes of 28 yeast cells across three conditions were generated and analyzed. 3D renderings of wild type, *Δncr1* and *Δnpc2* cells (n = 14, 6, 8 cells respectively) were used to measure the volume and pairwise distances between LDs and vacuoles. Together, we found 596 droplet meshes in 74 membrane meshes (i.e., 245 LDs in wild-type cells, 208 in *Δncr1* cells and143 in *Δnpc2* cells, respectively; Figure 5A). Together, 56 central vacuoles could be segmented, and in those cells, we found 272 LDs. Both, *Δncr1*, and *Δnpc2* cells had significantly larger LDs compared to the wild type as seen in Figure 5A. A Mann-Whitney U-Tests showed the significance of these results by giving p-values of 0.0037 and 0.0002 for *Δncr1*, and *Δnpc2* cells, respectively. Furthermore, our method allows for studying ingested LDs and LDs outside of vacuoles separately. LDs inside vacuoles are color-coded in purple in Figure 5B together with the key statistics for each droplet object. This includes a number for each droplet, the distance to the vacuolar membrane in µm, the droplet volume in µm^3^ and the percentage of its area in contact with the vacuole. Accordingly, LDs consumed by the vacuole have 100% of their surface area in contact with the vacuole, while those attaching on the outside have variable percentages, i.e., 22.43% for the green droplet in the inset of Figure 5B. LDs outside of the vacuole have of course 0% of their surface in contact and a given distance >0 from the vacuole membrane. This analysis allows for determining contact formation between both organelles in 3D in a fully automated manner. It reveals that there is no statistically significant difference in droplet uptake between wild-type and *Δncr1*, or *Δnpc2* cells supporting the findings from visual inspection of X-ray reconstructions. When measuring droplet volumes for every condition in cells with intact vacuoles, we found that the LDs outside of the vacuole were considerably smaller than those inside the vacuole in *Δncr1*, and *Δnpc2* cells compared to wild-type cells (Figure 5C-E). This suggests a selective defect in intravacuolar lipid digestion in cells lacking functional NCR1 or NPC2. It has been shown that lipid hydrolysis can take place inside but also outside of vacuoles in yeast [52]. Thus, one can speculate that LDs destined for cytosolic hydrolysis are compositionally and functionally different from those ingested into the vacuole, and that NPC proteins are important for lipid mobilization specifically from intravacuolar LDs. Together, our pipeline can identify subtle lipophagic phenotypes between wild type and NPC-deficient yeast cells, which could not be identified without automated organelle segmentation of X-ray images and subsequent statistical analysis.

**Figure 5.**
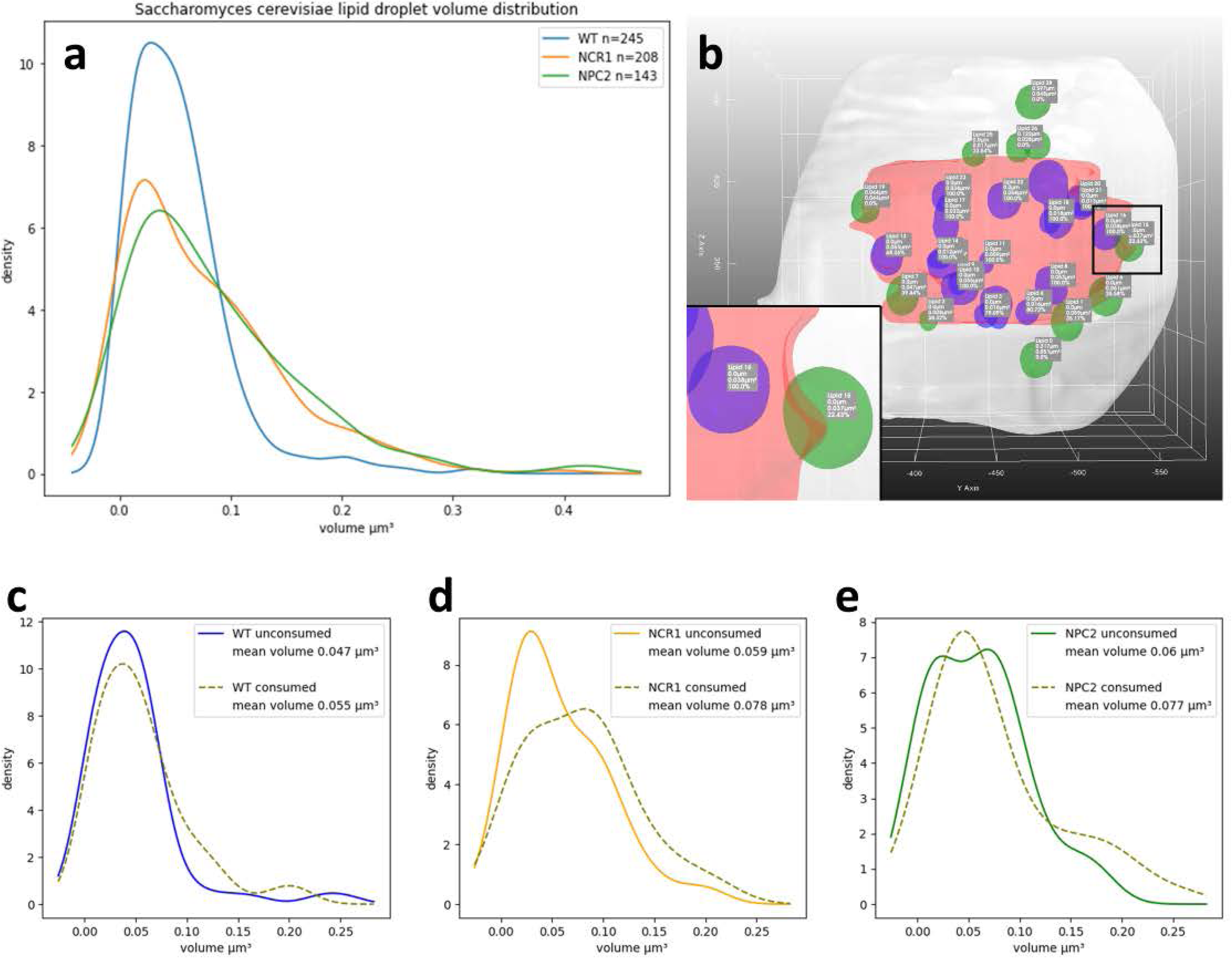
Quantitative analysis of lipophagy in yeast cells from segmentation CNN output. (**A**) Kernel density plot of droplet volume measured from 3D-rendered LDs inside and outside of vacuoles in wild-type cells (blue line), *Δncr1* cells (orange line) and *Δnpc2* cells (green line). Model output with LDs inside vacuoles color-coded in pink, and LDs outside of vacuoles in green (**B**). Labels are provided showing statistics for each droplet. The inset in (**B**) corresponding to the black box show one droplet inside of the vacuole in purple, and another one attaching in green. Identified LDs were separated into those inside vacuoles (‘unconsumed’) and inside vacuoles (‘consumed’), and their respective volume was calculated for wild-type cells (**C**), *Δncr1* cells (**D**) and *Δnpc2* cells (**E**) and plotted as kernel density plot. See main text for further details.

## Discussion

In this work, we developed a deep learning approach to automatize the analysis of fluorescence and X-ray microscopy images of yeast during lipophagy. This form of autophagy is increasingly recognized as important regulatory process for maintenance of cellular energy homeostasis. Dysfunction of lipophagy has been shown to play an important role in development of obesity, development of type 2 diabetes and oxidative stress as well as in protein aggregation during neurodegeneration [53, 54]. Lipophagy is often studied in yeast, Saccharomyces cerevisiae, because this model organism is straightforward to manipulate genetically and also because the yeast vacuole is large enough to be amenable to light microscopic investigation. This allows for counting the number of LDs inside vacuoles as a measure for lipophagy under various starvation conditions and genetic backgrounds [7, 14, 45, 55, 56]. While reliable and largely reproduceable between different human observers, visual annotation of vacuole phenotypes is tedious. Given the recent progress in machine learning, in particular of deep learning in microscopy [57], we chose to implement a neural network for automatizing this task. In contrast to classical image analysis, CNNs can automatically optimize feature learning and extraction for phenotype classification by learning from annotated examples (so-called supervised learning). This approach has been used extensively for phenotype classification in yeast, including analysis of subcellular protein localization, cell morphology, cell budding, genetic interactions and autophagy based on light microscopy image data [9, 58, 59, 60]. We show that accumulation of lipids in the vacuole can be reliably distinguished from lipid-poor vacuoles in which FM4-64 staining concentrated in the limiting membrane, giving the vacuolar labeling a ring-like morphology. Lipid-filled vacuoles are often accompanied by bright intensity of BODIPY 495/503 in LDs, and we find that our classification CNN can reliably distinguish both phenotypes (Figure 1). This enables us to show that yeast lacking functional NCR1 or NPC2 have a defect in clearance of vacuolar lipids in stationary phase lipophagy (Figure 1) [8]. Our classification CNN complements existing methods for automated, deep learning-based phenotype scoring in yeast cells. For example, multi-layer CNNs have been used for classifying subcellular localizations of green-fluorescent-protein-tagged proteins in yeast, and classification of such autophagy markers in the vacuole [9, 58, 61, 62].

Since diffraction-limited fluorescence microscopy lacks the resolution to study the vacuolar phenotype at the ultrastructural level, we employed a combination of fluorescence and X-ray microscopy and found that cells lacking functional NCR1 or NPC2 deposit more intravacuolar lipid material which is in line with their function as lipid transporters out of this organelle (Figure 2A) [8, 10, 11, 63]. Vacuoles and LDs can be unequivocally identified by their characteristic shape and X-ray absorption but also by the overlap in fluorescence from organelle-specific markers, emphasizing the potential of correlative fluorescence and X-ray imaging (Figure 2B). We make use of another and more complex CNN to segment these organelles from X-ray reconstructions. CNNs have been used extensively for segmentation of computational tomography data in the clinical imaging communities [33, 34, 35, 36]. However, few studies have used CNNs for segmentation of SXT data of individual cells [25, 26, 64]. Using an U-net-type architecture for the segmentation task and a traditional, watershed-based approach for the subsequent instance segmentation, we keep the computational burden and need for training data at a minimum and are nevertheless able to automatically segment vacuoles and LDs from X-ray image stacks and thereby to quantify the extent of droplet uptake into the vacuole. Our model thereby differs from previous attempts to segment organelles from X-ray tomogram reconstructions, as we use partially fluorescence annotated X-ray data for training [25, 26, 64].

We find that *Δncr1*, and *Δnpc2* cells can internalize LDs into the vacuole similar to control cells, however, the intraluminal LDs are enlarged in cells lacking functional NCR1 and NPC2 (Figure 3). 3D rendering allows us not only to determine the extent of lipophagy but also to determine the surface fraction of the segmented LDs in contact with vacuoles. This is important, as it enables us to study contact sites between organelles in a fully automated manner from SXT data. SXT has previously been shown to allow for studying membrane contact sites between lysosomes and mitochondria and between insulin granules and mitochondria but only on a qualitative level or upon tedious manual segmentation [65, 66]. Our quantitative analysis allows for determining the contact surface between LDs and vacuoles in 3D from X-ray images in an automated manner, which will be important for analysis of contact sites between other organelles in the future. We find that LDs inside vacuoles were larger for *Δncr1*, and *Δnpc2* cells compared to wild-type cells, while the size of droplets outside the vacuole is comparable (Figure 5C-E). This result suggests that specifically intravacuolar lipid mobilization depends on NPC-proteins in yeast. Interestingly, cytosolic hydrolysis of stored triacylglycerols and ergosterol esters in the cytosol has been shown to be coupled to a functional vacuole, and the products of cytosolic digestion of LDs were found to be efficiently used for membrane remodeling during growth resumption [52]. It will be interesting to study, whether droplet utilization under growth resumption is differently affected by lack of NCR1 or NPC2 compared to stationary phase lipophagy.

A recent study used freeze-etch EM to study lipophagy in cells lacking functional NCR1 or NPC2 during nitrogen starvation and found evidence for impaired engulfment of LDs into the vacuole [7]. This was accompanied by reduced formation of vacuole domains in this study. In additon, autophagic vesicles inside the vacuole were found to be smaller in NPC-deficient yeast compared to wild-type cells [7]. We cannot confirm these findings, as droplet docking to the vacuole and engulfement of LDs does not seem to be the affected step in *Δncr1*, and *Δnpc2* cells in our experiments. Rather, we find that the droplet volume is larger, which would indicate that cells lacking NPC proteins cannot properly degrade the ingested LDs. This conclusion is also in line with our fluorescence imaging data and points to a defect in hydrolysis coupled lipid export from the vacuolar compartment. Normal droplet ingestion into vacuoles based on a biochemical assay (i.e., vacuolar degradation of Erg6 associated with LDs) was also found in another recent study [67]. Similarly, vacuolar domains, which are thought to be essential for droplet docking during lipophagy appear to be normal in *Δncr1* and *Δnpc2* cells during stationary phase and glucose starvation [44, 67]. Since the mode of induction of lipophagy seems to affect the molecular machinery, which yeast cells use to execute lipophagy [2], it is possible that NPC proteins are differently involved in stationary phase compared to other forms of lipophagy. In line with this hypothesis are recent findings, that lipid imbalance by disturbed PC synthesis and ER stress affect vacuole domain formation in an NCR1- and NPC2-dependent manner during lipophagy [67].

While LDs are rich in ergosterol and yeast NPC proteins are important for sterol export from the vacuole, NCR1 and NPC2 might transport other lipids as well. In fact, we showed recently, that yeast NPC2 binds a variety of phospholipids and can adapt its binding pocket to the different lipid ligands [63]. If this is similarly the case for NCR1, one could hypothesize, that both proteins are needed to shuttle a variety of lipids out of the vacuole during starvation. As a consequence, absence of either protein should slow intravacuolar lipid digestion, simply because hydrolysis products would not be exported from the vacuole and thereby could inhibit further lipid cleavage by product feedback inhibition. We are currently testing this hypothesis using the tools described in this study. We also found that vacuoles of *Δncr1*, and *Δnpc2* cells sometimes appear fragmented, suggesting that vacuole fusion is compromised in a subpopulation of NPC-deficient cells (Figure 2D and E). This could be a consequence of defective ergosterol transport into the vacuolar membrane, as ergosterol is needed for vacuole fusion [8, 68]. Future studies are needed to unravel the molecular mechanisms underlying the function of NPC proteins in various forms of lipophagy, and the tools provided here will be instrumental for this endeavor.

## Materials and Methods

### Yeast strains, labeling and preparation for microscopy

The WT BY4741 yeast strain was obtained from the Euroscarf culture collection, while the BY4742 ncr1D::KanMX4 and BY4742 npc2D::KanMX4 strains were kindly provided by Prof. Christopher Beh of Simon Fraser University, Burnaby, Canada. The cells were grown in YPD media consisting of D(+)-glucose monohydrate (Merck, 1.08342.1000), 2% Bacto Peptone (BD Chemicals, 211677), 1% yeast extract (Merck, 1.03753.0500), and 0.02% adenine (Sigma-Aldrich, A-2786). Prior to experimentation, the cells were cultured in 5 mL of media in a 50 mL Falcon tube at 30°C with 150 rpm until they reached the stationary phase. The optical density at 600 nm (OD600) was then measured using a WPA Biowave CO8000 (Biochrom Ltd, UK), and the cells were washed three times with sterile water before being diluted to a final OD600 concentration of 0.1. The diluted cells were incubated for 22 hours under anaerobic conditions in YPD media containing 0.1 µg/mL *N*-(3-Triethylammoniumpropyl)-4-(6-(4-(Diethylamino) Phenyl) Hexatrienyl) Pyridinium Dibromide (FM4-64, Thermo Fisher, T3166) and 0.1% Tween 80 (Sigma-Aldrich, P4780) with 5 µg/mL DHE (Sigma-Aldrich, 810253P). After 22 hours, the yeast was washed three times with PBS, resuspended in the original growth media, and incubated under aerobic conditions for 24 hours. For imaging, the cells were diluted in PBS to a final OD600 concentration of 1.0 and labeled with 0.5 mg/mL 4,4-Difluoro-1,3,5,7,8-Pentamethyl-4-Bora-3a,4a-Diaza-*s*-Indacene (BODIPY 493/503; Thermo Fisher, D3922), a marker for LDs, for 3 minutes.

### Fluorescence microscopy and soft X-ray tomography

All fluorescence images used for vacuole phenotyping by deep learning were acquired using a Leica DMIRBE microscope with a 63x 1.4 NA with an oil immersion objective. Images were obtained using a neutral density filter. For FM4-64 dye a standard rhodamine filter set with 535 nm (50nm bandpass) excitation filter, 565 nm dichromatic mirror and 610 nm (75 nm bandpass) emission filter was used. BODIPY 493/503 was imaged using a standard fluorescein filter set, with a 470 nm (20nm bandpass) excitation filter, 510 nm dichromatic mirror and a 537 nm (23 nm bandpass) emission filter. The fluorescent sterol, DHE, was imaged using a special design filter cube in the UV range with a 335 nm (20 nm bandpass) excitation filter, 365 nm dichromatic mirror and 405 nm (40 nm bandpass) emission filter. For SXT, cells were treated as described above, seeded on the poly-D-lysine coated R2/2 grids (Quanti-foil, 100 Holy Carbon Films, Grids: HZB-2 Au), and a small volume of 270 nm gold beads is added to the grids as fiducial markers. The sample is plunge-frozen and stored in liquid nitrogen before imaging. The imaging of the grids is done on the U41-PGM1 TXM beamline over a tilt angle range of 120-125° with 1° tilt steps using an X-ray photon energy of 510 eV and a 25-nm zone plate. Before SXT, fluorescence images of the samples were taken in the same set up under cryogenic conditions using a Zeiss LD EC Epiplan Neofluar 100x objective with NA=0.75 and standard rhodamine and fluorescein filter sets. The image pixel size is 9.8 nm for SXT and 148 nm for fluorescence microscopy.

### Registration of X-ray tomograms, 3D image reconstruction and alignment with fluorescence

For the alignment, the Bsoft software [68] was used with the gold beads as fiducial markers [69]. Individual frames of tomograms with motion blur were manually removed before processing. After the alignment, tomograms were binned by a factor of two. The reconstruction was carried out in Tomo3D using the filtered back projection algorithm [70]. For correlation of fluorescence and X-ray images, sum projections of portions of reconstructed 3D X-ray stacks were calculated to match the depth of field of the fluorescence microscope. Fluorescence images were deconvolved as described above prior to alignment with the reconstructed SXT sum projections.

### Fluorescence image preprocessing and cell segmentation using the circular Hough transform

Fluorescence images of DHE, FM4-64 and BODIPY 493/503 were first deconvolved using a Richardson-Lucy algorithm with 30 iterations and a theoretical point spread function using the Deconvolution Lab software in ImageJ as described [71]. Image stacks of DHE were first corrected for residual autofluorescence by subtracting the last from the first image in each stack, as described [72]. For isolating individual cells, the bright field image, acquired in parallel, was used. One image has a size of 512×512 pixels, with an average of 70 yeast cells, and for easier cell segmentation, shading correction was required. For that, a copy of each BF was blurred with a Gaussian filter with standard deviation of 20 pixels and subtracted from the original bright field image. This procedure was automatized as ImageJ Macro and retained all cellular features, while removing background intensity variations. Automated detection and segmentation of yeast cells is an important step for large-scale live-cell imaging experiments. It can be demanding, if cells tend to cluster, bud extensively or growth in heterogeneous environments, such as microfluidic devices, and a variety of advanced algorithms including CNNs have been developed recently to address this problem, e.g., [61, 73, 74, 75, 76, 77, 78, 79, 80]. Since our experimental setup is much simpler with well separated cells on a rather homogeneous background, we can rely on a morphometric segmentation procedure, the circular Hough transform, which detects objects based on their circularity. The algorithm was used as Macro-callable ImageJ plugin (https://imagej.net/Hough_Circle_Transform developed by Dr. Benjamin Smith, UCB Vision Science, Berkeley, USA). We implemented this plugin in a Macro script after intensity thresholding to generate a binary image and applying a Sobel edge filter to get an approximate outline of the cell periphery. The plugin determines centroid positions of each yeast cell defined as circular object using a simple geometric transformation from polar to cartesian coordinates. The determined centroid positions are then used to extract sub-images of size 32×32 pixels for each cell in an ImageJ Macro script. The entire procedure of shading correction of bright field images and cell segmentation using the circular Hough transform is illustrated in Fig. S1.

### Deep learning model setup for vacuole phenotype classification

After generating the single yeast cell images, they must be annotated to be utilized for training and validation. Each of the 32×32 pixel images is manually annotated into two categories: ring-like vacuole corresponding to the ‘healthy’ phenotype and ‘sick’ phenotype based on lipid-filled or fragmented vacuoles as visible in the FM4-64 vacuole staining. This step of the procedure is crucial, as wrongly annotated images can influence the output of the deep learning model. A third category was added for vacuole morphologies, which did not match either category (‘unknown’). Therefore, each of the images, which is used for training and validating the model, is carefully annotated, and divided into these three categories. After annotation, the images are converted from 32-bit image (as this is output from deconvolution) to 8-bit images, and the two categories are split into two groups: training and validation. The CNN for fluorescence phenotype classification was implemented in Python with Tensorflow (https://www.tensorflow.org/) and Keras (https://keras.io/). Input images are the 32×32 pixel 8-bit greyscale images for the vacuole marker, which were normalized by dividing with 255. To increase the variability in our training data without the need for acquiring new data, we performed the following augmentations in Keras: random rotation, where the same image is rotated by different angles, horizontal flipping, translations using a width and height shift of 0.2 pixels, and random zoom where the new image is a zoom of a part in the original data. This step is done to prevent the model from overfitting and thus improving the ability to generalize. Each input image is passed through convolution blocks, the first block includes a 2D convolution layer and a Max-pooling layer. The activation function used in this setup is ReLU for ensuring non-linearity. After the last layer, a flattening layer is included, which converts the output from the last layer to a 1D array. Here, a dropout of 0.5 is used to prevent overfitting. The 1D array is subsequently fed into a dense layer, which consists of two sublayers. The first part of the dense layer has an activation function ReLU with 128 units. The output layer, which is the second dense sublayer, has a sigmoid activation function, used for binary classification. As loss function the binary cross entropy is used. The ‘Adam’ optimizer is then used to adjust and update the weights in the network during training. Epochs were set for 200, and as a validation of the model, the loss and accuracy are measured through the entire process. The number of images for training, validation and testing are given in Tab. 1. A ROC curve is calculated after running the model on the validation data. The model is then used on unseen data from both conditions, and it calculates the percentage of ring-like and lipid-filled vacuole phenotypes for both cases.

### Generation of training data for the segmentation model, preprocessing and data augmentation

Training data for the segmentation CNN was created by hand using ImageJ and 3D slicer. Many good segmentations of unseen data were added after post-processing (see below). Training data was verified using correlative fluorescent markers when available, and otherwise overlaid on the original data and evaluated by experts. In the final version of the training set, the cell membrane and droplet channel have 539 slices of training data across 28 tomograms. The vacuole channel has 56 slices across 21 tomograms. Each image in the Z-stack is rescaled to 512×512 using bicubic interpolation. The original dimensions are stored in the program memory, such that the original pixel sizes can be recovered, after processing. The pre-processing consists of resizing to 512×512 pixels and quantile normalization using a gaussian distribution with 0 mean and 1 standard deviation. This ensures consistent input for data that can have considerably different distributions. The procedure for this quantile normalization is described in the following pseudocode below:

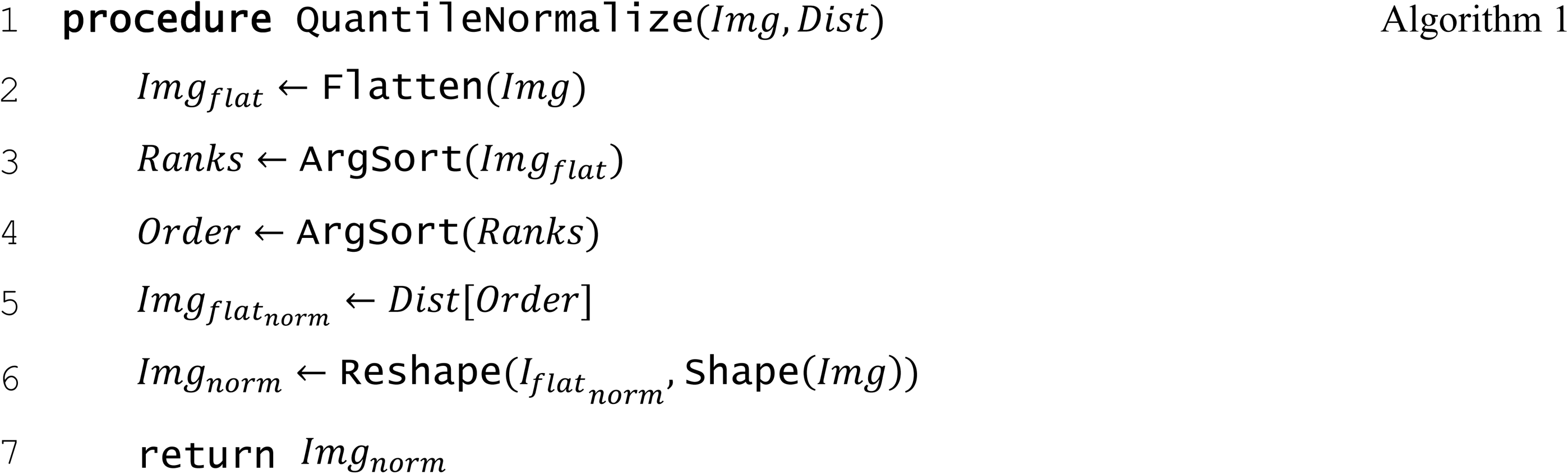

Image augmentation includes 90-, 180-, and 270-degree rotations and elastic deformations. The rotations are multiples of 90, such that no information is lost to black bars or zoom. Elastic deformations are implemented using Simard et al.’s algorithm. α and σ parameters are set to 430 and 20, respectively, and were selected to get a meaningful distortion while maintaining recognizable features and patterns. A random seed is generated prior to deformation, such that all five images receive the same distortion.

### Architecture and training of the segmentation model

We decided to use an U-Net based architecture as already proven to be powerful for similar image processing tasks [33, 34]. Other U-Net style architectures were tested but were dismissed to keep the complexity and capacity for overfitting down. The architecture is implemented using five encoder blocks, a bridge, five decoder blocks, and a single convolutional layer with sigmoid activation. Each encoder and corresponding decoder are connected through concatenation. Each successive encoder doubles the number of filters (starting with 32) and each decoder halves it again. Encoder blocks contains 2 sets of convolutions, batch normalization, and ReLU activation with the second set having half the number of filters in the convolutional layer. Each encoder is succeeded by a 2×2 max pooling layer, halving the feature size in each dimension. The Decoder block is an encoder block preceded by an up-sampling layer, which doubles the feature map size in each dimension. All convolutional layers use zero padding to keep the image dimensions consistent and in powers of two. The input shape is 512×512×5. Five consecutive images are used to aide in consistency, reduce the effect of noise from individual slices and give the network some 3D context to combat the missing wedge problem.

A model is trained for each class. The training data is categorized according to the biological sample / tomogram and for each ‘epoch’, a random training pair is sampled from each tomogram. This reduces bias towards certain samples and phenotypes with more training data available. The Adam optimizer was selected based on an empirical review of optimizers. A compound loss function consisting of Dice and Top K cross entropy was selected, based on [81, 82].

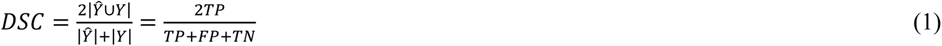

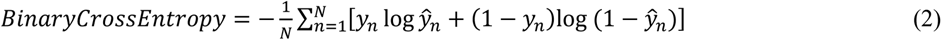

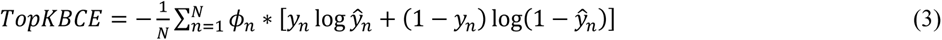

where 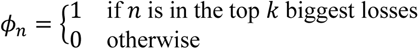

Models were trained iteratively on each channel, as more training data was created. The vacuole channel was trained using the cell membrane channel as initial model weights.

### Postprocessing and analysis of segmented 3D X-ray image stacks

Outlier removal removes artifacts outside of the cell membrane. This is accomplished by thresholding the mean Z-projection of the cell membrane prediction stack and expanding it using binary dilation. This mask is then applied to the cell membrane channel. The similar masking option uses the post-processed cell membrane segmentation as a mask for other channels with the masking option enabled. This prevents misclassifications of cellular components outside of the cell membrane (e.g., fiducial markers which may resemble LDs). The averaging option is implemented using a 3D gaussian filter which increases consistency in areas with low prediction confidence and slices causing unexpected drops resulting in false negatives. Hole filling is implemented using SciPy’s *binary_fill_holes* [83]. The implementation fills the holes by dilating the complementary of the input. Hole filling is conducted after instance segmentation, such that gaps between cells and cellular components are not wrongly filled. Hole filling can aid in predictions where smaller sections fall below the classification threshold and generate holes in the membrane or vacuole. The convex constraint is implemented by reducing point clouds to convex hulls [84]. This can help in cases where the segmentation is known to be convex but yields a concave shape. After mesh generation using marching cubes, meshes may be smoothed using Laplacian smoothing.

For the analysis, 28 *S. cerevisiae* yeast cell tomograms were segmented using cell membrane, vacuole, and droplet channels. 14 WT, 6 NCR1 and 8 NPC2. The segmentations were sorted into those where a vacuole could be identified and segmented and those that could not. For each droplet in all segmentations, volume, distance to nearest vacuole, whether it was consumed by a vacuole and the sample type were recorded. Distance was estimated by raytracing between centers of mass. LDs were considered consumed if their center of mass was enclosed by the nearest vacuole’s surface. The distribution of droplet volumes between wild type and NCR1 were compared. Additionally, in all samples with well-segmented vacuoles, the distributions of volumes of LDs inside and outside vacuoles were plotted against each other using gaussian kernel density estimation and p-values were acquired using method Mann-Whitney U-test and the Kolmogorov–Smirnov test. Similar comparisons were made under other conditions but did not yield p-values under 0.05.

Image simulations were carried out in ImageJ, while tomographic reconstruction of the simulated yeast phantom images was implemented in the skimage Python library skimage.transform.

## Abbreviations

BODIPY493/503: 4,4-Difluoro-1,3,5,7,8-Pentamethyl-4-Bora-3a,4a-Diaza-*s*-Indacene
CNN: convolutional neural network
DHE: dehydroergosterol
*Δncr1*: yeast deficient in NCR1
*Δnpc2*: yeast deficient in NPC2
DSC: Dice similarity coefficient
EM: electron microscopy
EVs: extracellular vesicles
FIB-SEM: focused ion beam milling scanning electron microscopy
FM4-64: *N*-(3-Triethylammoniumpropyl)-4-(6-(4-(Diethylamino) Phenyl) Hexatrienyl)-pyridinium dibromide
LDs: lipid droplets
NCR1: yeast homologue of Niemann Pick C1 protein
NPC: Niemann Pick type C
NPC2: Niemann Pick C2 protein
OD600: optical density at 600 nm
ReLU: rectifier linear unit
SXT: soft X-ray tomography
UV: ultraviolet
YPD: yeast extract peptone dextrose

## Acknowledgments

DW acknowledges funding from the Villum foundation (grant no. 35865) and from the Danish Research Council (grant ID: 2032-00139B).

## Data availability

The code for the classification and the segmentation CNN is available at https://github.com/Wuestner-Lab/LipoSeg. Selected examples of 3D reconstructions from X-ray tomograms will be made availability at https://zenodo.org with the link provided at the GitHub page.

## Supplemental figure legends

**Figure S1.** Circular Hough transform for segmentation of yeast cells from bright field images. Bright field images (**A**) were first flat-field corrected by subtracting a copied version blurred with a 20-pixel wide gaussian filter from the original image in 32-bit format (**B**). An edge filter was applied to enhance the cell border (**C**) followed by intensity thresholding and binarization (**D**). A circular Hough transform was applied providing the approximate radius and centroid position for each cell in the field of view (**E**). From that, individual cells could be identified (**F**), cropped and saved as separate image files for all four channels (not shown) as trainings and validation data for the classification CNN. This figure is related to figure 1.

**Figure S2.** Workflow of combined X-ray and fluorescence microscopy of yeast cells. Yeast cells were grown and optionally labeled with fluorescent markers of interest. Grids were coated with poly-D-lysine before cells were added to them and allowed to settle for approximately 25 min. Gold beads were added to the grids before they were plunge frozen, to serve as fiducial markers. The grid was placed in a special holder and kept under cryo-conditions during acquisitions of first fluorescence light images followed by SXT. Based on the gold beads, the raw tilt series were aligned with B-soft and subsequently 3D reconstructed with Tomo3D. TomoJ was used to manually align the tomograms with the fluorescence images. This figure is related to figure 2 to 4.

**Figure S3.** Comparison of manually and automated segmentation of X-ray image stacks. Dice similarity coefficient (‘DSC’) for model prediction versus ground truth as function of frame number along the z-axis for entire cells (black curves) or LDs (green curves) for a partially unseen X-ray stack (**A**) and for a fully unseen X-ray stack (**B**). The ground truth segmentations were obtained from hand-segmented image stacks, while the predicted segmentations are the output of the segmentation CNN model. Example frames for the ground truth versus predicted segmentations (**C**). This figure is related to Figure 4.

**Figure S4.** Illustration of the missing wedge problem using a synthetic yeast cell phantom. A yeast cell phantom was generated as 2D image with a visible cell membrane, vacuole and LDs and additive Gaussian noise (**A and C**). Using this phantom as input image 2D projections were calculated in parallel beam geometry as acquired on a real X-ray tomography system. This gives the Radon transform (also called sinogram) for either 180 degrees of projection angles (**B**) or 130 degrees (**D**), the latter corresponding to the SXT set up from the synchrotron BESSY II, used in this study. The 2D image of the yeast cell was reconstructed from both sinograms using a filtered back projection algorithm with a Hamming filter for regularization for the full range tomogram (**E**) or the limited-angle tomogram (**F**). The reconstruction artifacts are clearly visible for the reconstruction with 130 degrees. Note, that for real SXT tomograms, the yeast cells are much smaller relative to the beam and detector geometry making that the artifacts shown here are much less pronounced in the real data. Thus, this figure is only an illustration of the tomographic principle causing reconstruction artifacts. This figure is related to Figure 4.

**Figure S5.** Additional examples for lipid storage phenotypes in NPC-deficient yeast cells. (**A and B**) Sum projections along the optical axis of five selected frames each from a SXT reconstruction are shown for *Δnpc2* cells to illustrate the lipid-filled vacuole including intraluminal vesicles (white arrows) and aberrant vacuole structure as well as formation of extracellular vesicles (blue arrow and zoomed box in **A**). (**C and D**) Side-views of 3D rendering obtained using the segmentation CNN for that cell shows that all LDs are ingested into the vacuole. The vacuole shape is a slightly deformed pseudo-hexagonal structure as inferred from the sum projections as well as from the 3D reconstructions. A zoomed version of panel (**C**) is shown in (**D**). Individual frames of the sum projection throughout the SXT reconstruction of the *Δnpc2* cell shown in the main text (**E**). Selected frames of a reconstructed stack (first four panels) and a sum projection (most right panel) of *Δncr1* cell with lipid-filled vacuole (**F**). This figure is related to Figure 4.

**Supplemental video.** Left panel: reconstruction of X-ray tomogram is shown along the z-axis; middle panel: predicted instances of cells (blue), vacuoles (red) and droplets (green); right panel: overlay of X-ray images segmentation output.

