## Supplemental figures for "Automated quantification of lipophagy in Saccharomyces cerevisiae from fluorescence and cryo-soft X-ray microscopy data using deep learning"

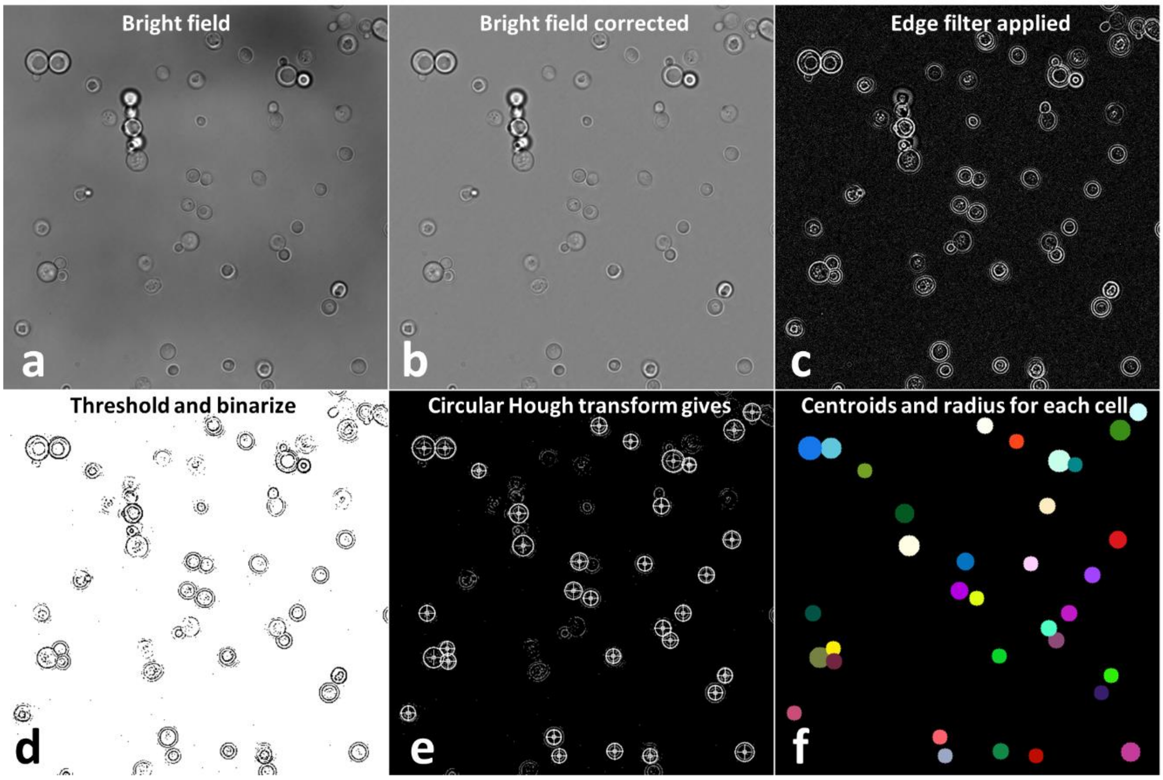


**Figure S1.** Circular Hough transform for segmentation of yeast cells from bright field images. Bright field images (**A**) were first flat-field corrected by subtracting a copied version blurred with a 20-pixel wide gaussian filter from the original image in 32-bit format (**B**). An edge filter was applied to enhance the cell border (**C**) followed by intensity thresholding and binarization (**D**). A circular Hough transform was applied providing the approximate radius and centroid position for each cell in the field of view (**E**). From that, individual cells could be identified (**F**), cropped and saved as separate image files for all four channels (not shown) as trainings and validation data for the classification CNN. This figure is related to figure 1.

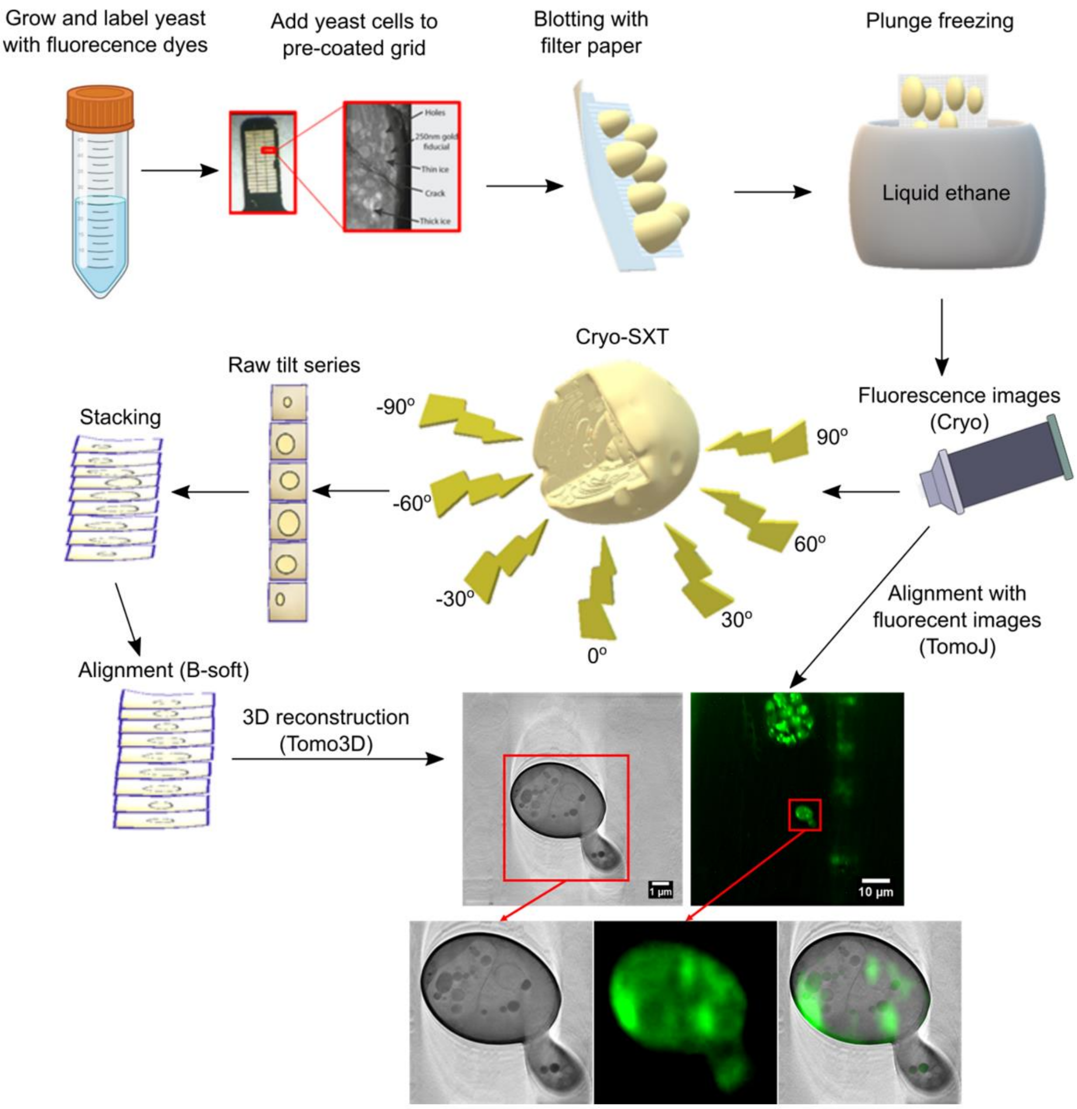


**Figure S2.** Workflow of combined X-ray and fluorescence microscopy of yeast cells. Yeast cells were grown and optionally labeled with fluorescent markers of interest. Grids were coated with poly-D-lysine before cells were added to them and allowed to settle for approximately 25 min. Gold beads were added to the grids before they were plunge frozen, to serve as fiducial markers. The grid was placed in a special holder and kept under cryo-conditions during acquisitions of first fluorescence light images followed by SXT. Based on the gold beads, the raw tilt series were aligned with B-soft and subsequently 3D reconstructed with Tomo3D. TomoJ was used to manually align the tomograms with the fluorescence images. This figure is related to figure 2 to 4.

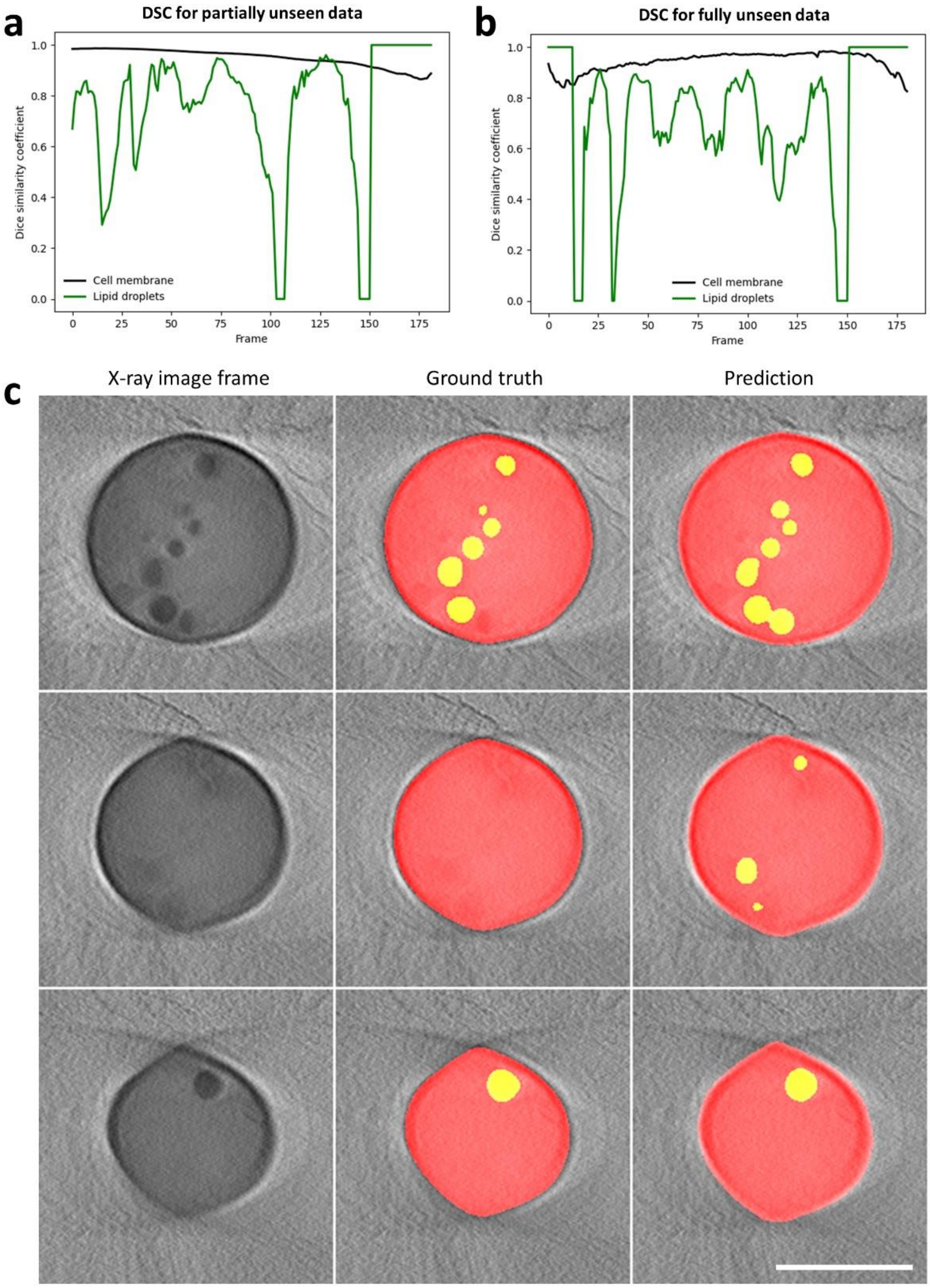


**Figure S3.** Comparison of manually and automated segmentation of X-ray image stacks. Dice similarity coefficient ('DSC') for model prediction versus ground truth as function of frame number along the z-axis for entire cells (black curves) or LDs (green curves) for a partially unseen X-ray stack (**A**) and for a fully unseen X-ray stack (**B**). The ground truth segmentations were obtained from hand-segmented image stacks, while the predicted segmentations are the output of the segmentation CNN model. Example frames for the ground truth versus predicted segmentations (**C**). This figure is related to Figure 4.

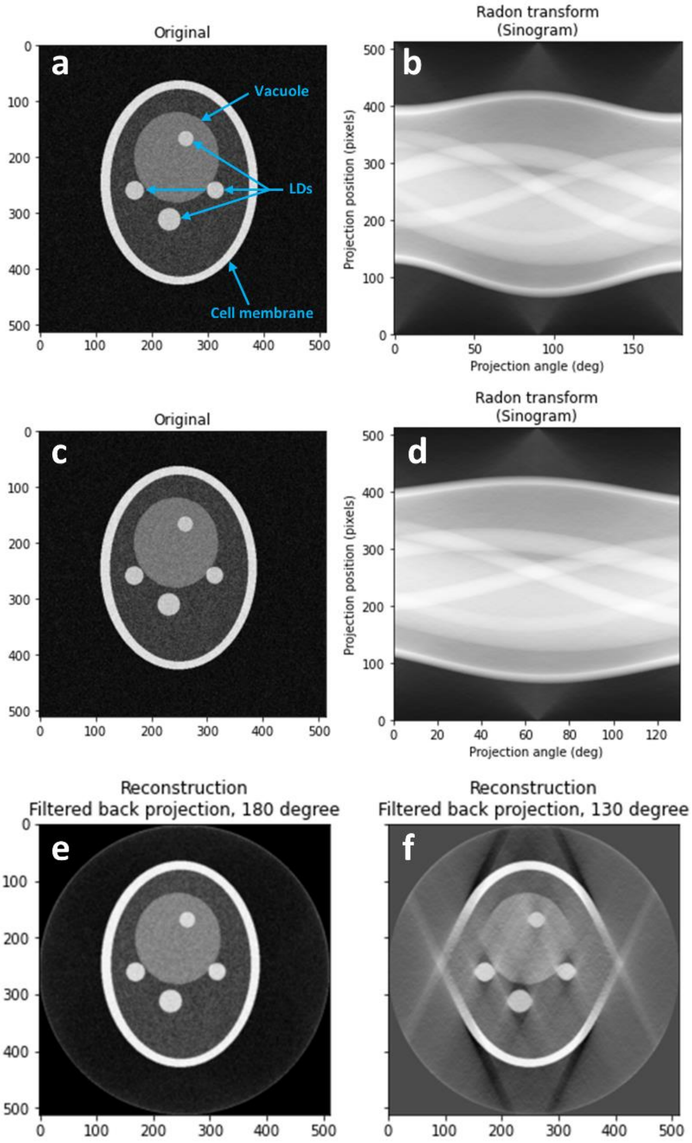


**Figure S4.** Illustration of the missing wedge problem using a synthetic yeast cell phantom. A yeast cell phantom was generated as 2D image with a visible cell membrane, vacuole and LDs and additive Gaussian noise (**A and C**). Using this phantom as input image 2D projections were calculated in parallel beam geometry as acquired on a real X-ray tomography system. This gives the Radon transform (also called sinogram) for either 180 degrees of projection angles (**B**) or 130 degrees (**D**), the latter corresponding to the SXT set up from the synchrotron BESSY II, used in this study. The 2D image of the yeast cell was reconstructed from both sinograms using a filtered back projection algorithm with a Hamming filter for regularization for the full range tomogram (**E**) or the limited-angle tomogram (**F**). The reconstruction artifacts are clearly visible for the reconstruction with 130 degrees. Note, that for real SXT tomograms, the yeast cells are much smaller relative to the beam and detector geometry making that the artifacts shown here are much less pronounced in the real data. Thus, this figure is only an illustration of the tomographic principle causing reconstruction artifacts. This figure is related to Figure 4.

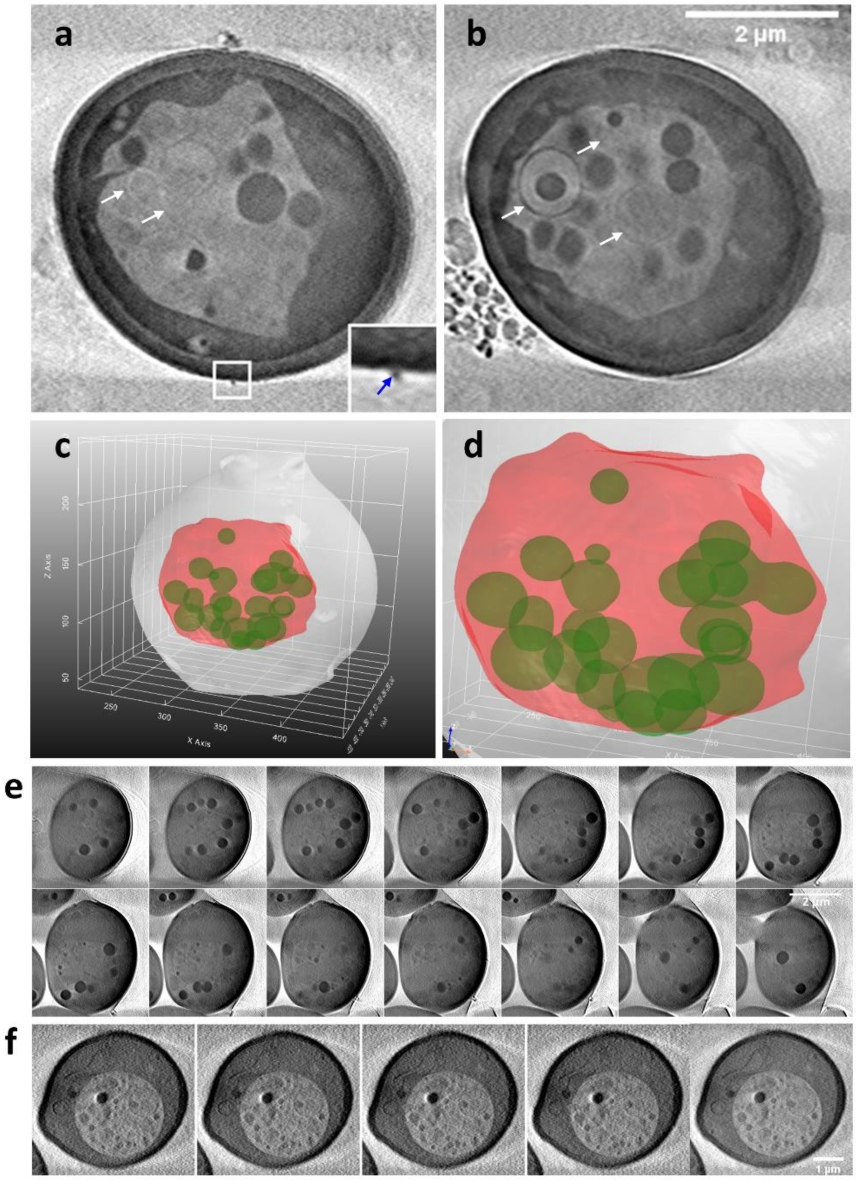


**Figure S5.** Additional examples for lipid storage phenotypes in NPC-deficient yeast cells. (**A and B**) Sum projections along the optical axis of five selected frames each from a SXT reconstruction are shown for *Δnpc2* cells to illustrate the lipid-filled vacuole including intraluminal vesicles (white arrows) and aberrant vacuole structure as well as formation of extracellular vesicles (blue arrow and zoomed box in **A**). (**C and D**) Side-views of 3D rendering obtained using the segmentation CNN for that cell shows that all LDs are ingested into the vacuole. The vacuole shape is a slightly deformed pseudo-hexagonal structure as inferred from the sum projections as well as from the 3D reconstructions. A zoomed version of panel (**C**) is shown in (**D**). Individual frames of the sum projection throughout the SXT reconstruction of the *Δnpc2* cell shown in the main text (**E**). Selected frames of a reconstructed stack (first four panels) and a sum projection (most right panel) of *Δncr1* cell with lipid-filled vacuole (**F**). This figure is related to Figure 4.
